# The influence of river discharge on gravel bar hyporheic microbial community structure and putative metabolic functions

**DOI:** 10.1101/2022.10.11.511717

**Authors:** Arnelyn D. Doloiras-Laraño, Joeselle M. Serrana, Shinji Takahashi, Yasuhiro Takemon, Kozo Watanabe

## Abstract

1. Microbial communities in the hyporheic zone (HZ) are important in self-purification as the riverbed is metabolically active and responsible for the retention, storage and mineralisation of organic matter transported to the surface water. However, studies exploring HZ microbial community responses to disturbances (e.g. floods) remain scarce.
2. Here, we characterised the microbial community structure among the three (downwelling, upwelling and intermediate) HZ points within and among gravel bars at high and low discharge levels in a dam-regulated river using 16S rRNA metabarcoding.
3. We observed significant dissimilarity in the microbial community at low discharge exhibiting local adaptation due to gravel bar spatial environmental heterogeneity. Moreover, the homogenisation effect resulted in similar microbial community structures among the three points within the gravel bars at high discharge. Microbial communities across adjacent gravel bars were dissimilar, potentially attributing to different bar morphologies.
4. Our study highlights the role of spatial environmental heterogeneity in the biological processes that govern microbial community structure at three hyporheic points in gravel bars at two discharge levels.
5. Our results are essential to understand the HZ microbial communities’ response to the river discharge levels.

## Introduction

In riverine ecosystems, the hyporheic zone (HZ) in gravel bars, where the surface and groundwaters meet, plays an essential role in providing habitat and refuge for microbial communities (Boano et al., 2014; Boulton & Stanley 1995) and in maintaining healthy waterways. In the HZ, heterogeneous environmental conditions can occur along the water pass from the downwelling zone (surface water movement toward the groundwater) at the bar head to the upwelling zone (groundwater movement toward the surface water) at the bar tail of the gravel bar water flows into the HZ (Hendricks 1993). This hyporheic flow carries different materials with it and other microbial communities, removing water pollutants and natural solutes.

Previous studies used molecular approaches to investigate how the hyporheic flow path influences microbial communities in HZs in gravel bars. For example, Lowell et al., (2009) demonstrated, using polymerase chain reaction-denaturing gel electrophoresis (PCR-DGGE), that microbial communities were spatially diverse along a 100-m hyporheic flow path with large environmental heterogeneity such as chemicals and nutrients. Kim & Lee (2019) studied how vertical flow directions (i.e. upwelling/downwelling) affect the microbial community using pyrosequencing and identified different microbial community compositions between the upwelling/downwelling zones.

As rivers are typically dynamic environments, understanding the community structure-related changes from pre- to post-flooding has long been an important research topic in disturbance ecology (Graham et al., 2021). However, previous studies focused only on surface water organisms, reporting behaviours such as refuge to the HZs during flooding (Miyake et al., 2020, Calderon et al., 2017) and community succession after disturbance. To the best of our knowledge, no study has focused yet on the disturbance ecology of HZ communities. Increased discharge in a river, such as a flood, can increase hydraulic pressure on the riverbed, potentially increasing the downwelling flow from the surface water to the HZ (Boano et al., 2014) or the flow velocity of the normally slow-flowing hyporheic water. Such changes in the hydrologic conditions could potentially result in HZ community disturbance during flooding. If a heterogeneous microbial community structure is formed along the HZ path of gravel bar under low discharge conditions, a disturbance occurring under high river surface water discharge, such as a flood, could homogenise it. To test this hypothesis, it is necessary to compare the HZ community divergence levels within the gravel bars between high and low discharge conditions.

Understanding the functional trait of each species could contribute to deepening our knowledge of the biological processes and ecological mechanisms that shape community structure variation along environmental gradients. Recent studies on freshwater microbial communities also estimated functional traits based on taxonomic annotation using databases (e.g. SILVA rRNA, Greengenes, National Center for Biotechnology, and Ribosomal Database Project) and 16S rRNA metabarcoding (Serrana et al., 2022; Galand et al., 2018; Fasching et al., 2020) Several studies have demonstrated the validity and utility of this approach, although the estimated functional traits were putative as 16S rRNA metabarcoding could not identify functional genes in the genome, and database limitations allowed annotating only a proportion of the taxa. However, applied cases are still scarce and further verifications would be required.

This study aimed to assess how river discharge influences the microbial community dynamics of the gravel bar HZs in a river under dam regulation. In particular, we aimed to (1) profile the microbial diversity and community structure among HZs at three hyporheic, i.e. downwelling, upwelling and intermediate, points of gravel bars, (2) identify the microbial community structure-influencing environmental factors among the HZ points, (3) compare the divergence levels of the microbial communities among the HZ points under low and high discharge conditions of the river surface water, and (4) identify the putative metabolic functions of the microbial communities on these sites under different discharge levels. Taken together, this study can provide valuable insights into the role of spatial environmental heterogeneity in the biological processes and ecological mechanisms that govern the local adaptation of microbial communities at taxonomic and functional levels in gravel bar HZs.

## Methods

### Study site and water sample collection

The Tenryu River is located in central Honshu, Japan, with a length and basin area of 213 km and 5,090 km^2^, respectively. The river originates from Lake Suwa in Nagano Prefecture and is discharged into the Pacific Ocean (Figure 1). The Funagira Dam (34°53’26’N, 137°48’54’) was the last major dam to be completed on the Tenryu River supplying water to the nearby Funagira Hydroelectric Power Station. The dam releases different water discharge levels throughout the year, thereby altering the water flow discharge of the river.

**Figure 1.**
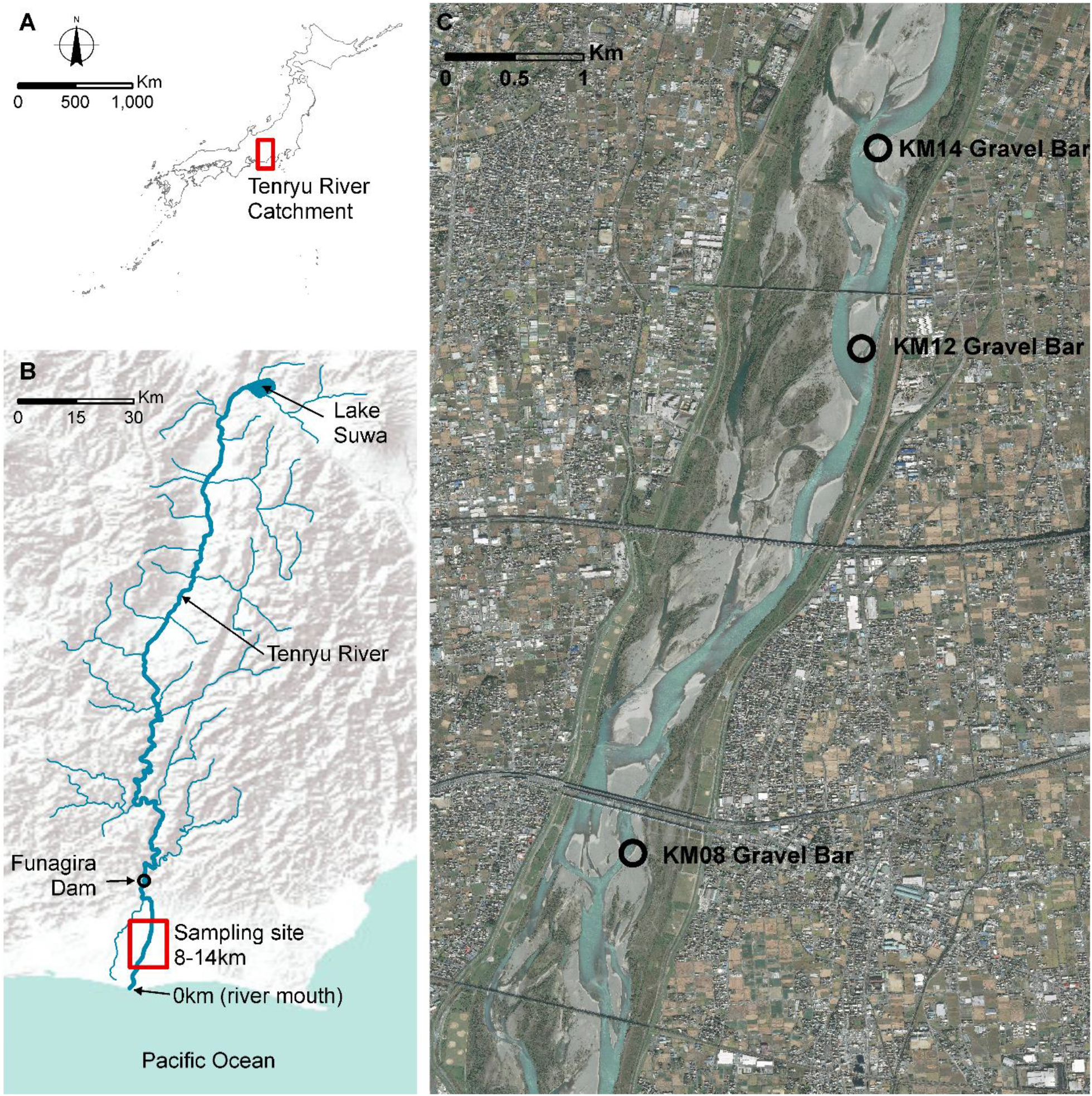
Detailed map of the sampling sites (A) The top panel shows the contour of the map of Japan indicating the Tenryu River. (B) The figure shows the location of Lake Suwa where the river originates and Funagira Dam over Esri elevation model. (C) Google map aerial photo of the three gravel bars (KM08, KM12 and KM14) from the river mouth.

Field surveys were conducted between November 13–17, 2019. The mean dam discharge was high (250 m^3^/s) and low (150 m^3^/s) between November 13–15 and 16–17, 2019, respectively. During the field survey, flow fluctuations reduced the water level to 80 cm, thereby changing gravel bar morphology. Three gravel bars located 8, 12, and 14 km (KM08, KM12 and KM14, respectively) from the river mouth were selected in this study. These three large gravel bars are distributed adjacent to each other within 6 km of the middle reach of the Tenryu River, with relatively similar environmental conditions. We collected and measured water samples and environmental parameters at three sampling points among each gravel bar site, i.e. the downwelling, upwelling and intermediate points. However, no sample was collected from the intermediate point of gravel bar KM14 as no hyporheic water was collected 20 cm from the ground. In total, we assessed 16 sampling points, *n* = 8 both at low and high discharges, in this study and collected water samples in triplicates for each sampling point (a total of 48 water samples).

A 1,220-mm-long solid steel cone piezometer (AXEL, Japan) was used to collect water samples from a depth of 20 cm from the ground surface at each hyporheic point. Hyporheic water samples of 250–500 ml were filtered using a 50-ml sterile syringe (Terumo) and 0.22-μm Sterivex™ filters (Merck Millipore, Merck KgaA, Darmstadt, 346 Germany) on-site. A negative field control sample was produced by filtering 1 l of ddH2O through a Sterivex™ filter, similar to the field samples. The filtered samples were preserved in 99% molecular-grade ethanol, brought to the laboratory, and stored at −20°C until processing for molecular analysis.

Water quality parameters, i.e. pH, electrical conductivity (EC), water temperature and turbidity, were recorded *in situ*. pH, EC, and water temperature were measured with a multi-parameter water quality checker (HORIBA, D-54, Japan). Turbidity was measured with a portable turbidity metre (TB-31, DKK-TOA, Japan). Suspended solids (SS) were collected by filtering 0.5 litres of hyporheic river water, passing it through a 1-mm sieve onto a GF/F filter (pore size: 0.7 μm), drying (105°C, 4h) and weighing on an electronic balance. Anions (F-, Cl-, NO_3_-N, SO_4_2-) were measured using ion chromatography (Metrohm, Basic IC plus 883, Swiss). Inorganic elements (Mg, Al, and Na) were measured using an inductively coupled plasma mass spectrometer (Agilent Technologies, Agilent8800 ICP-QQQ, Japan).

### DNA extraction, library preparation and amplicon sequencing

Total genomic DNA was extracted from each filter using the Qiagen PowerSoil DNA Isolation Kit (Qiagen, Hilden, Germany) following the manufacturer’s protocol. DNA quality and concentration were determined using a NanoDrop spectrophotometer (Thermo Scientific Nanodrop 2000), and the QuantiFluor dsDNA System (Promega, Madison, USA). The amplicon library was prepared through a one-step PCR protocol using modified primer sequences targeting the V4-V5 hypervariable region of the 16S SSU rRNA gene from the Earth Microbiome Protocol (Caporaso, et al., 2012). The PCR was performed using a T100 Thermal Cycler (Bio-Rad Laboratories, USA) and Phusion high-fidelity DNA polymerase (New England, Biolabs) for the PCR amplification. The 15-μl PCR reaction mixture consisted of 3 μl of 5X Phusion GC Buffer, 0.5 μl each of both the forward and reverse primers (10 μM), 0.6 μl dNTPs (2.5 mM), 0.45 μl DMSO, 0.3 μl Phusion Polymerase (1U), and 1 μl of template DNA at a concentration of 5 ng/μl. The PCR cycling conditions were as follows: initial denaturation at 98°C for 3 min, 25 cycles of denaturation, annealing, and extension at 98°C for 15 s, 67°C for 30 s, and 72°C for 30 s, respectively, followed by a final extension cycle at 72°C for 5 min. The amplicon size was 420 bp. Two negative controls i.e. field control sample and nuclease-free water, were included in the experiment to monitor the potential contamination from DNA extraction to PCR amplification. A total of 50 amplicon libraries (i.e. 2 negative controls, and triplicates of 16 points) were constructed. The PCR products were verified on a 2% agarose gel using gel electrophoresis. Next, we quantified the concentration of each PCR product using the KAPPA Illumina Library qPCR Quantification kit (Kappa Biosystems, Wilmington, MA, USA). Equimolar concentrations of each amplicon library were then pooled. The pooled samples were purified and size-selected using solid-phase reversible immobilisation beads (Beckman Coulter, Inc. CA). We used the Agilent Bioanalyzer 2100 system to assess the quality of the pooled library by the obtention of a single clear band of 420 bp. Sequencing was conducted on an Illumina Miseq (Illumina, Inc. San Diego, CA, USA) using a v3 Miseq sequencing kit (300 × 2, MS-102-3003) with 30% PhiX (Illumina, Inc. San Diego, CA, USA) and a starting concentration of 4nM denatured to a final concentration of 8pM.

### Read processing and taxonomic assignment

Raw paired-end reads were verified for quality using FastQC v0.11.8 (Andrews 2010). The raw reads were demultiplexed using the QIIME v2018.11 package (Boylen et al., 2019). Demultiplexed sequence data were then quality screened, processed and denoised using the DADA2 pipeline v1.16 package (Callahan et al., 2016). The reads were quality-filtered and truncated into 100-bp fragments. The chimeric sequences and singletons were also removed. Based on the read error files, the reverse reads displayed poor read quality. Therefore, only the forward reads were processed for downstream analyses. Amplicon sequence variants (ASVs) were inferred from the sequence data using the DADA2 pipeline v1.16 package. Taxonomic ASV identification was performed against the SILVA SSU database v132 using the SILVA ACT (www.arb-silva.de/aligner) (Pruesse et al., 2012). Subsequent analyses were performed at the ASV-level taxonomic unit.

### Statistical analyses and visualisation

All statistical analyses and visualisations were performed using the R software v4.0.1 (R Core, 2021). Read number normalisations were performed in each sample using median sequencing depth before the analyses. Species diversity at each point (alpha diversity) was determined using Chao1 richness (Chao 1984), Shannon diversity (Shannon & Weaver 1949), and Simpson diversity (Simpson 1949) via the plot_richness command in the phyloseq package (McMurdie & Holmes 2014). The mean alpha diversity values for each replicate in the HZ points were computed. Analysis of variance (ANOVA) was conducted via phyloseq to test whether the mean alpha diversities were significantly different among the three or two points per gravel bars. Microbial communities between points of gravel bars (beta diversity) at ASV-level taxonomic units were computed using Bray–Curtis dissimilarity (Bray & Curtis 1957) using the distance function in the vegan package, and visualised by non-metric multidimensional scaling (NMDS) using the plot_ordination function in the phyloseq package (Dixon, 2003). Permutational multivariate analysis of variance (PERMANOVA) was measured using the phyloseq package to test potential significant differences between points per gravel bars. Distance-based redundancy analysis (dbRDA) was performed to determine the environmental factors influencing the microbial community structure among the hyporheic points and visualised via the cca function of the vegan package. We identified the putative metabolic functions using the functional annotations for the prokaryotic (FAPROTAX) v1.2.3 (Louca & Doepeli 2017) database. The mantel test (Legendre & Legendre 2012) was conducted via the mantel.correlog function with 9.999 permutations to test for significant correlations between beta diversity based on taxonomic structure among the points and that based on the putative metabolic function structure.

## Results

### Microbial diversity and community composition

We generated a total of 15,346,131 raw reads from the 50 amplicon libraries. The reads one and six of the two negative controls were removed for subsequent analyses. After quality filtering, chimeric sequence removal and denoising, we retained a total of 5,439,168 reads (Table S1). Out of the inferred 19,592 ASVs, only 5,453 ASVs exhibited genus level, and from the latter, 260 ASVs species-level assignments. At the genus level, *Flavobacterium* (440,130 reads, 21% of the total number of reads assigned to genera) was the most abundant among the three different points followed by *Methylobacter* (115,342 reads, 6%), *Sediminibacterium* (92,722 reads, 5%), and *Rhodoferax* (88,531 reads, 4%) (Supplementary Figure 1C). Proteobacteria was the most abundant phylum (1,321,906 reads, 33% of total numbers of reads assigned to phyla), followed by Bacteriodiota (1,311,523 reads, 32%), Verrucomicrobiota (269,793 reads, 7%), Acidobacteria (217,836 reads, 6%), and Actinobacteria (60,563 reads, 2%) (Supplementary Figure 1D).

In each gravel bar, the NMDS ordination (Figure 2A) showed clustering of the three hyporheic points within the gravel bars at low discharge while such clustering could not be observed at high discharge using ASV-level taxonomic units (Figure 2C). PERMANOVA (Table 2) showed significant dissimilarity in the community structure among the three points at low discharge (*p* = 0.004) but no significant difference at high discharge (*p* = 0.326). The NMDS results (Figure 2B and 2D) of the three gravel bars showed clustered patterns under both high and low discharge conditions. PERMANOVA revealed significant community divergence among the three gravel bars at low (*p* = 0.005) and high (*p* = 0.001) discharge (Table 3).

**Figure 2.**
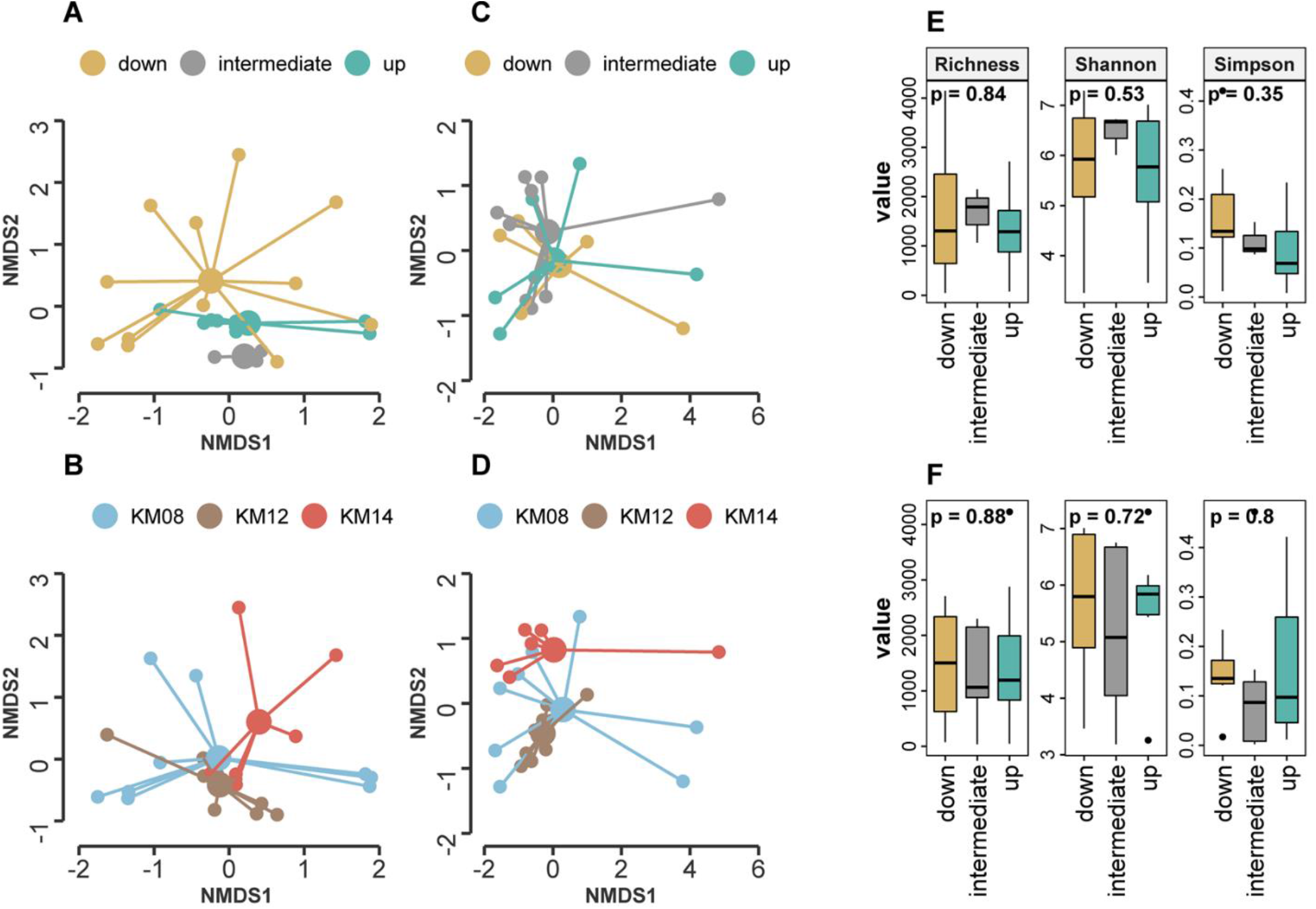
Non-multidimensional scaling (NMDS) showing the microbial community structure in (A) low discharge by hyporheic point (B) low discharge by gravel bar (C) high discharge by hyporheic point (D) high discharge by gravel bar. Box plot showing the alpha diversity per hyporheic point (i.e. downwelling, upwelling and intermediate) (E) Alpha diversity in low discharge (F) Alpha diversity in high discharge.

The alpha diversity chao1 richness, Shannon, and Simpson indices ranged between 35– 4023, 3.18–7.33, and 0.002–0.471, respectively, among the 16 points based on the ASV-level taxonomic units (Table S2). We observed no significant difference between the alpha metrics between the points within gravel bars under both low and high discharge conditions (Figure 2E and 2F).

The dbRDA results (Figure 3A and 3B) showed that turbidity (*p*=0.002), EC (*p*=0.001), and Al (*p*= 0.019) significantly influenced the microbial community structure as environmental factors at low discharge, while pH (*p*=0.033) and Ca (*p*= 0.011) were the environmental factors influencing microbial communities at high discharge (Table 1).

**Figure 3.**
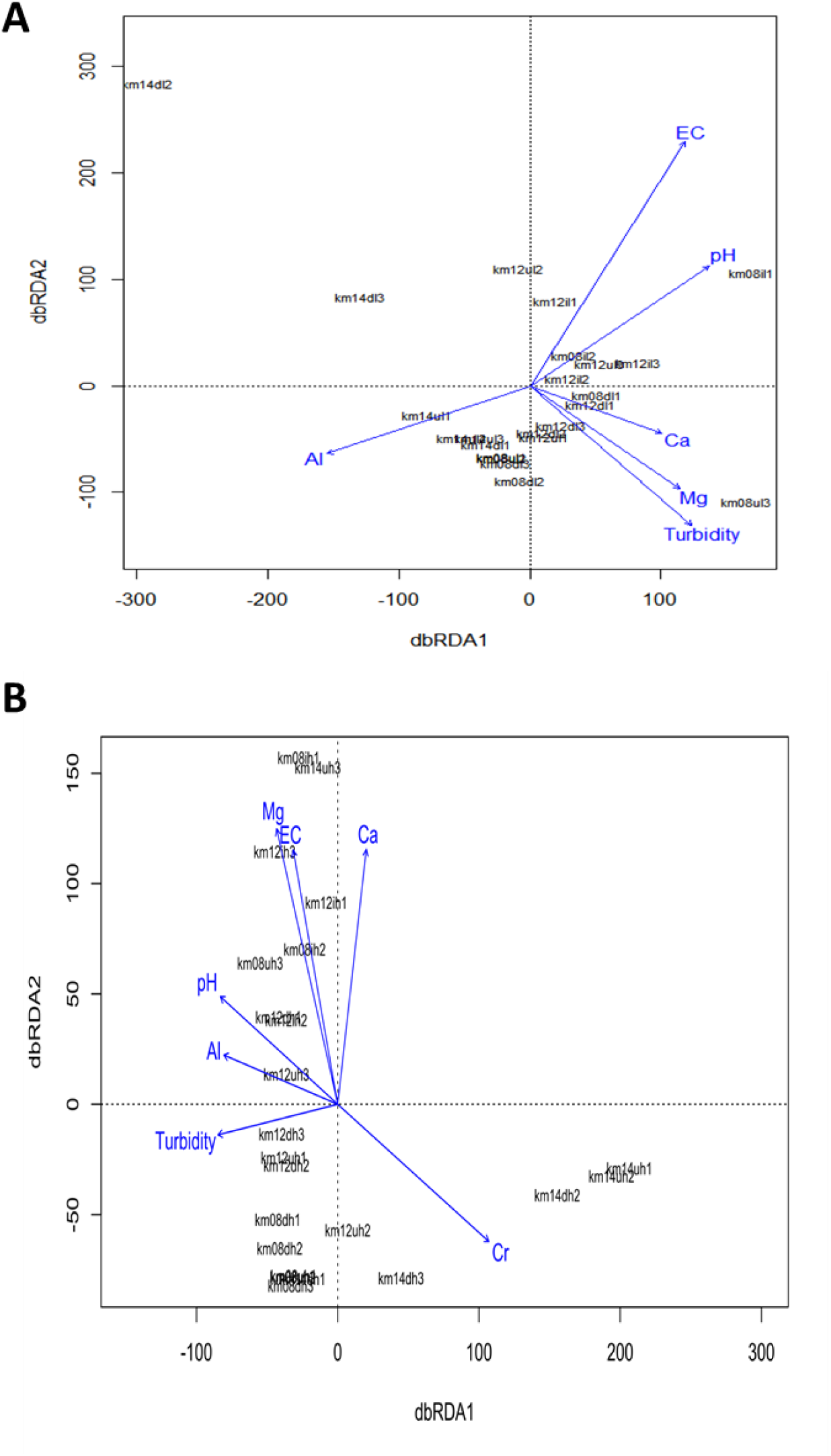
The dbRDA plot showing the significant environmental factors influencing microbial community: (A) dbRDA in low discharge (B) dbRDA in high discharge.

**Table 1.**
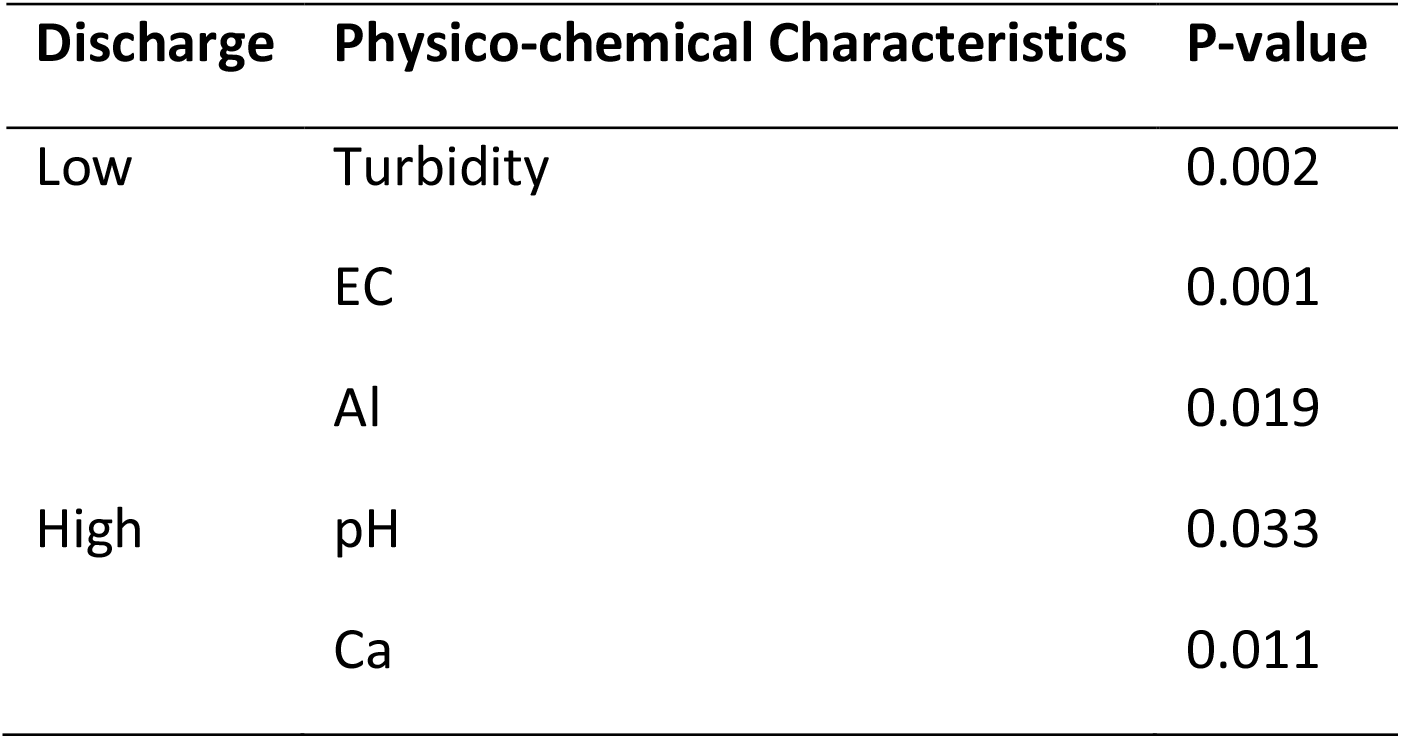
Distance-based redundancy (dbRDA) analysis shows the environmental factors affecting the microbial community structure for each gravel bar in two discharges levels.

**Table 2.** PERMANOVA of the three points in low and high discharge.

| Discharge | Df | SumsofSqs | MeanSqs | F Model | R2 | Pr(>F) |
| --- | --- | --- | --- | --- | --- | --- |
| Low | 2 | 1.1201 | 0.56006 | 2.0685 | 0.16458 | 0.004 |
| High | 2 | 0.6663 | 0.33314 | 1.0589 | 0.09575 | 0.326 |

**Table 3.** PERMANOVA of the three gravel bars in low and high discharge.

| Discharge | Df | SumsofSqs | MeanSqs | F Model | R2 | Pr(>F) |
| --- | --- | --- | --- | --- | --- | --- |
| Low | 2 | 1.0656 | 0.53279 | 1.9491 | 0.15657 | 0.005 |
| High | 2 | 0.4433 | 0.72166 | 2.6171 | 0.2073 | 0.001 |

### Putative metabolic function of the hyporheic microbial communities

A total of 2,962 ASVs (15.12%) out of 5,453 ASVs were assigned to at least one putative metabolic function. We identified 67 out of 92 putative metabolic functions in the FAPROTAX database. Chemoheterotrophy and aerobic chemoheterotrophy obtained annotated putative metabolic function throughout the 16 points (Figure S2). Methanotrophy and methylotrophy were described as dominant in the downwelling points while methanogenesis was dominant in the upwelling points both at low and high discharge. Aerobic ammonia oxidation was the most abundant putative metabolic function identified at the intermediate point. We also described a significant positive correlation between beta diversity based on community structure and putative metabolic function structure (Figure 4).

**Figure 4.**
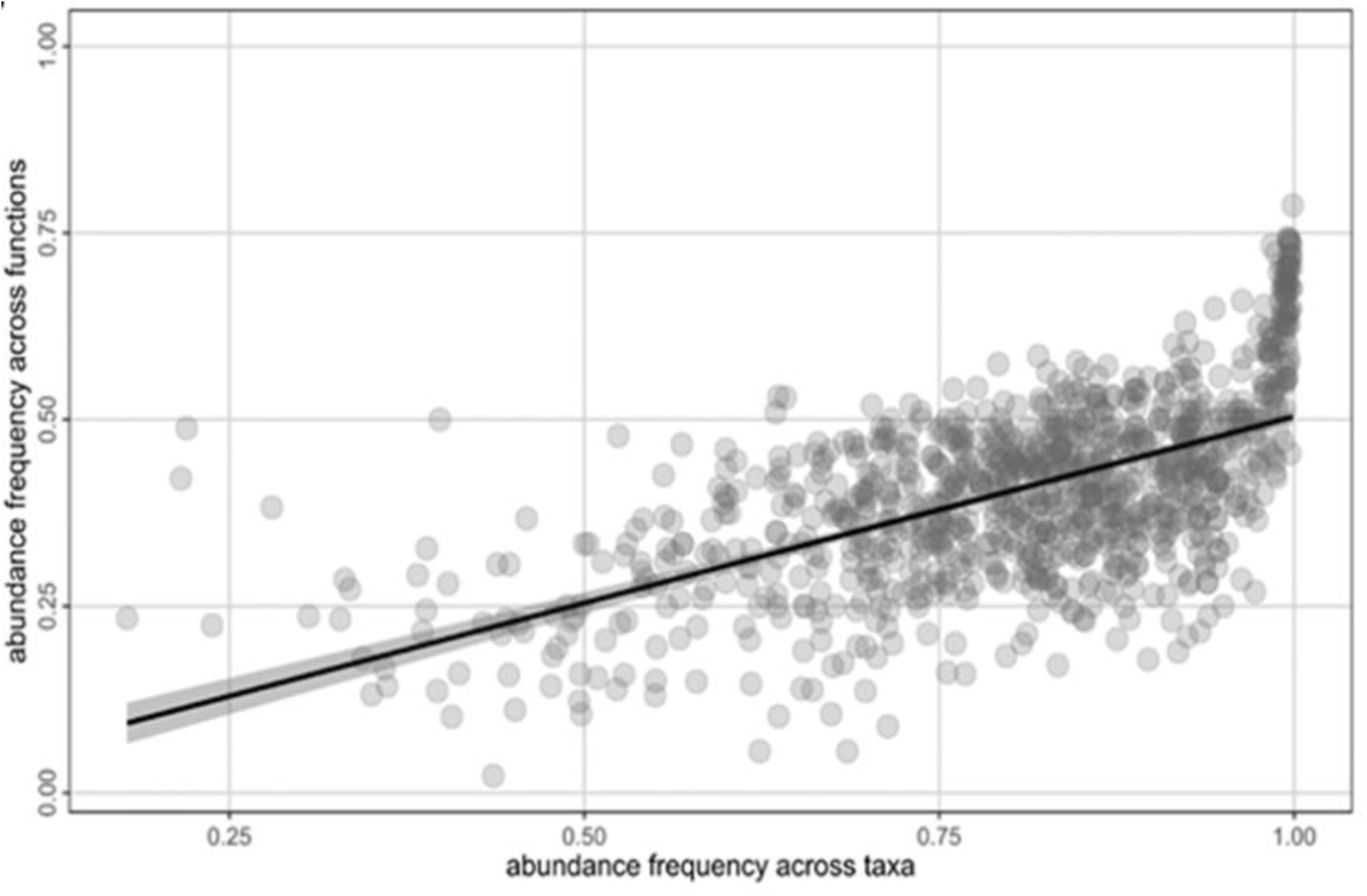
Mantel test analysis of abundance frequency based on taxa β-diversity versus abundance frequency based on putative metabolic functions using the Bray-Curtis index.

## Discussion

This study aimed to characterise and contrast the microbial community structure and putative metabolic function among the hyporheic i.e. downwelling, upwelling and intermediate points within and among gravel bars between two (i.e. low and high) discharge levels of a dam-regulated river. The community divergence is defined as the set of sampling units that becomes more dissimilar in composition so that pairwise dissimilarities increase over time (Buckley et al.,2019; Buckely et al., 2021). PERMANOVA revealed community patterns such as divergence or convergence by testing the significant compositional differences among the sample groups. Our results revealed significant microbial community divergence among the three points within the gravel bars under low discharge conditions. This could be attributed to local adaptation reflecting environmental heterogeneity among hyporheic points within the gravel bar at low discharge.

We demonstrated that turbidity, EC, Al, pH, and Ca were the environmental factors that significantly shaped microbial community divergence. Certain microbial community groups cannot withstand when metal concentrations increase in the environment. Moreover, metal concentrations play an essential role in influencing microbial communities (Zeglin 2015). For example, Feris et al., (2003) described a significant relationship between streambed metal concentrations in HZ and microbial community structure in six different rivers. Turbidity is essential in water microbial communities as it influences the processes within the primary producers, such as phytoplankton and photosynthetic bacteria. These primary producers rely on light energy and oxygen production for growth and variable water chemical properties, that in turn, influence bacterial nutrient use in the river ecosystem (Wagner et al., 2015).

A significant finding underlines the similarity of microbial community structure among the three hyporheic points observed under high discharge conditions. These results support our hypothesis that an increased river surface water discharge would also represent mean hydrologic changes in HZs. The disturbance among the HZ microbial communities might rise during the high discharge condition introducing homogenisation. This could be explained by the potential mechanism of mixing hyporheic water between the three points. However, determining the driving process was not possible since the surface water was not examined. Understanding how communities respond to disturbances such as a flood is essential to identify biological and ecological processes that determine their assembly and predict future effects on diversity and function (Marmonier et al., 2012). To the best of our knowledge, ours is the first study to investigate how increased river water discharge influences heterogeneous microbial communities among the hyporheic flow path in gravel bars. The results of this study provided clear evidence that HZ is not effective as a refuge from increased river water discharge as discussed in previous studies in invertebrates (Boulton & Stanley, 1995; Dole-Olivier et al., 1997).

The contrasting results of community divergence observed between two spatial scales, i.e. within and between gravel bar scales, provided valuable insights into the role of environmental heterogeneity in microbial community structure in the HZ at different spatial scales. We observed significant community divergence between 2-km distance gravel bars that remained even under high discharge conditions. However, we could not observe any significant divergence within the gravel bar scales under high discharge conditions. This divergence between gravel bars could be attributed to their bar geomorphology and sediment characterisation (Ock et al., 2015). Since microorganisms are adapted to the local environment of each gravel bar and the geographic distance between the bars was relevant, inter-bar migration that could homogenise the community structure among the bars was not evident regardless of the discharge level.

This study described ecologically important putative metabolic functions such as nitrification, denitrification and methanogenesis. These ecological functions were involved in biogeochemical processes in the HZ. We identified a dominant putative metabolic function at each hyporheic point (Supplementary Figure 2). Methanogenesis was abundant at the upwelling, while methanotrophy and methlylotrophy at the downwelling points in both discharge levels. This coincides with the results of Jones et al., (2015), stating that methanogenesis accounted for all the respiration in anoxic sediments and 0.3%–0.6% of the total respiration. These findings implied that HZ is essential in the carbon cycle such as methylotrophy, methanotrophy and methanogenesis in gravel bars. Moreover, the HZ through methanogenesis appears to be an important pathway for organic cycling and is a potential source of labile organic carbon on the surface of stream water. Putative metabolic functions, chemoheterotrophy and aerobic chemoheterotrophy, were the most dominant throughout all the 16 points, suggesting the importance of metabolically active microorganisms in the HZ. Mainly the abundant bacteria (e.g. Proteobacteria, Acidobacteria, and Verrucomicrobiota) contributed to chemoheterotrophy and aerobic chemoheterotrophy as putative metabolic functions. Our study also implies that HZ is suitable for the thriving of heterotrophic microorganisms.

Furthermore, we identified a positive correlation between taxonomic structure- and putative metabolic function structure-based beta diversity (Figure 5). This finding implies that the taxonomic structure divergence observed between points or gravel bars reflects different putative metabolic functions that emerged among the heterogeneous environments. This result is in good agreement with previous studies on stream microbial communities (Serrana et al., 2022; Galand et al., 2018; Fasching et al.,2020). However, it should be noted that the use of 16S rRNA gene sequences to predict metabolic function is constrained by the limited information content of the amplicon and limited databases. Hence, the putative metabolic function would require further validation using metagenomics and metatranscriptomics analyses that could provide comprehensive information on functional genes.

We observed no significant difference in alpha diversity among any of the three points in the three gravel bars under either low or high discharge conditions. This result coincided with those of Nelson, et al., (2019) and Sackett, et al., (2019), showing no significantly different variation level in microbial communities among the points in the gravel bar. We consider that each point can harbour similar alpha diversity levels with species turnover among the points. Potential mechanisms for the similar alpha diversity among the points, such as the environmental capacity for the number of species that each point can hold or limit the number of detectable species per point by metabarcoding, should be explored in future studies. Although alpha diversity did not change in hyporheic locations, we assume that microbial community structure might do, given the environmental heterogeneity of HZ in the gravel bar.

## Conclusions

In conclusion, spatial environmental heterogeneity among the HZ at low discharge resulted in microbial community structure divergence at three hyporheic (i.e. downwelling, upwelling and intermediate) points within the gravel bars. Our finding highlights how high discharge influences microbial community convergence as explained by the homogenisation in HZs. Putative metabolic functions were identified in the three hyporheic points, correlating with the taxonomic community structure and were determined to understand the process of microbial ecology dynamics in HZs. Furthermore, we identified environmental factors such as turbidity, EC and Al as drivers of divergence between hyporheic points within gravel bars. To further understand the biological mechanisms of microbial communities in the HZ, we believe that future studies should focus on metagenomics and metatranscriptomics to better understand the functional genes and activity of microbial communities in the HZ in gravel bars.

## Data availability statement

The raw sequences data was submitted to the National Center for Biotechnology Information (NCBI) Sequence Read Archive PRJNA855985.

## Authorship contribution statement

**Arnelyn D. Doloiras-Laraño:** Conceptualisation, Methodology, Investigation, Data curation, Formal analysis, Visualisation, Writing – original draft. **Joeselle M. Serrana:** Investigation, Formal analysis, Writing – review and editing. **Shinji Takahashi:** Investigation, Data curation, Methodology, Writing – review & editing. **Yasuhiro Takemon:** Investigation, Data curation, Writing - review & editing, Funding acquisition. **Kozo Watanabe:** Conceptualisation, Writing – review & editing, Resources, Supervision, Project administration, Funding acquisition.

## Declaration of Competing Interest

The authors declare no competing financial interest.

## ACKNOWLEDGEMENT

We acknowledge the substantial field work assistance of Dr. Maribet Gamboa, Somar Israel Fernando, and the Kyoto University Disaster Prevention Institute. We would also like to thank Dr. Naohito Tokunaga of Ehime University, Division of Analytical Biomedicine Advance Research Support Center (ADRES) and Dr. Paul Johnston of Leibniz-Institute of Freshwater Ecology and Inland Fisheries (IGB) for their assistance in the sequencing and data processing, and Micanaldo Ernesto Francisco for making a detailed map of the sampling sites. This work was supported by the Japan Society for the Promotion of Science (JSPS) Grant-in-Aid for Scientific Research (22H00571, 22H01627).

Appendix A. Supplementary data

## SUPPLEMENTARY TABLES AND FIGURES

**Supplementary Table 1.**
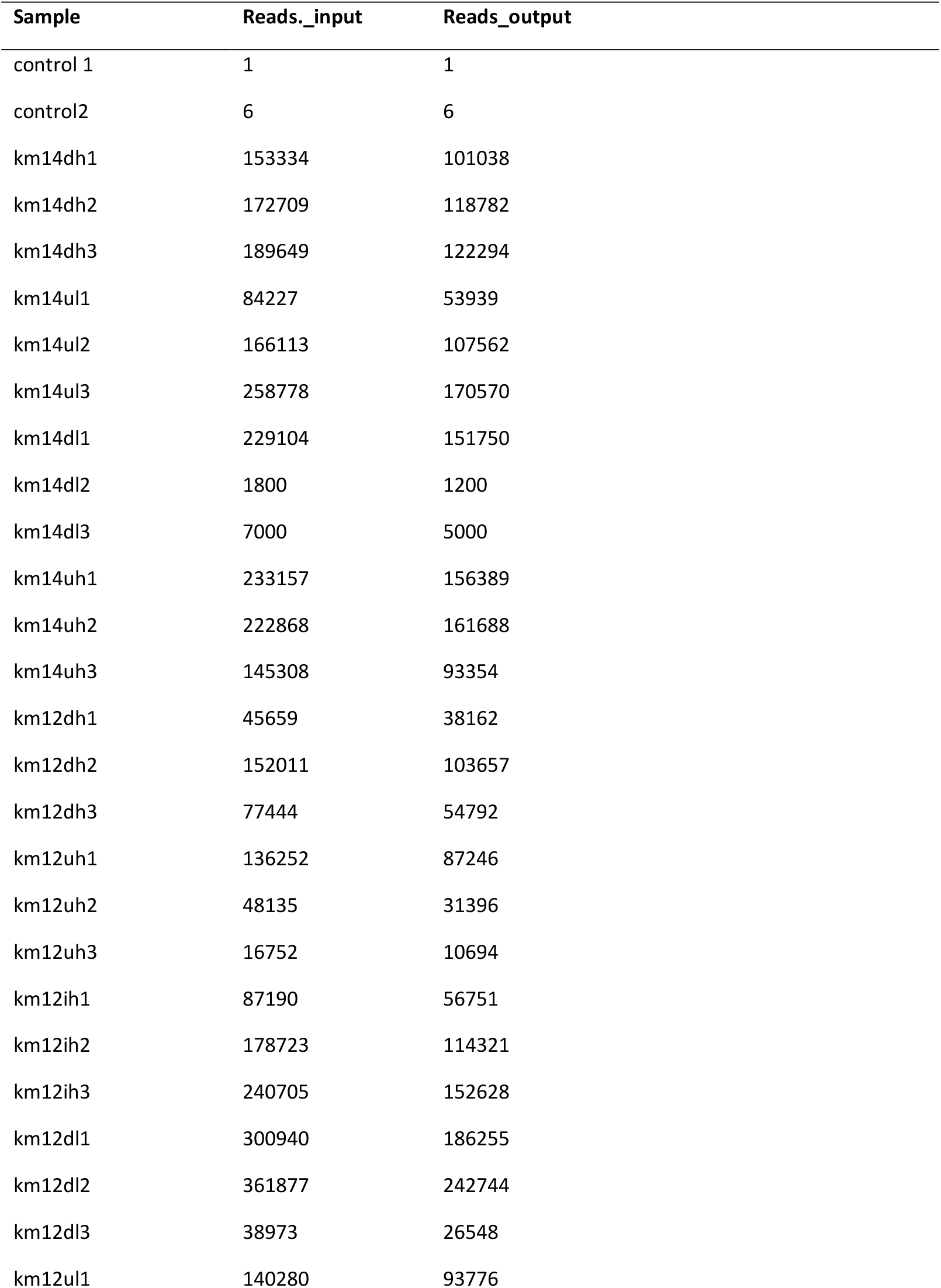
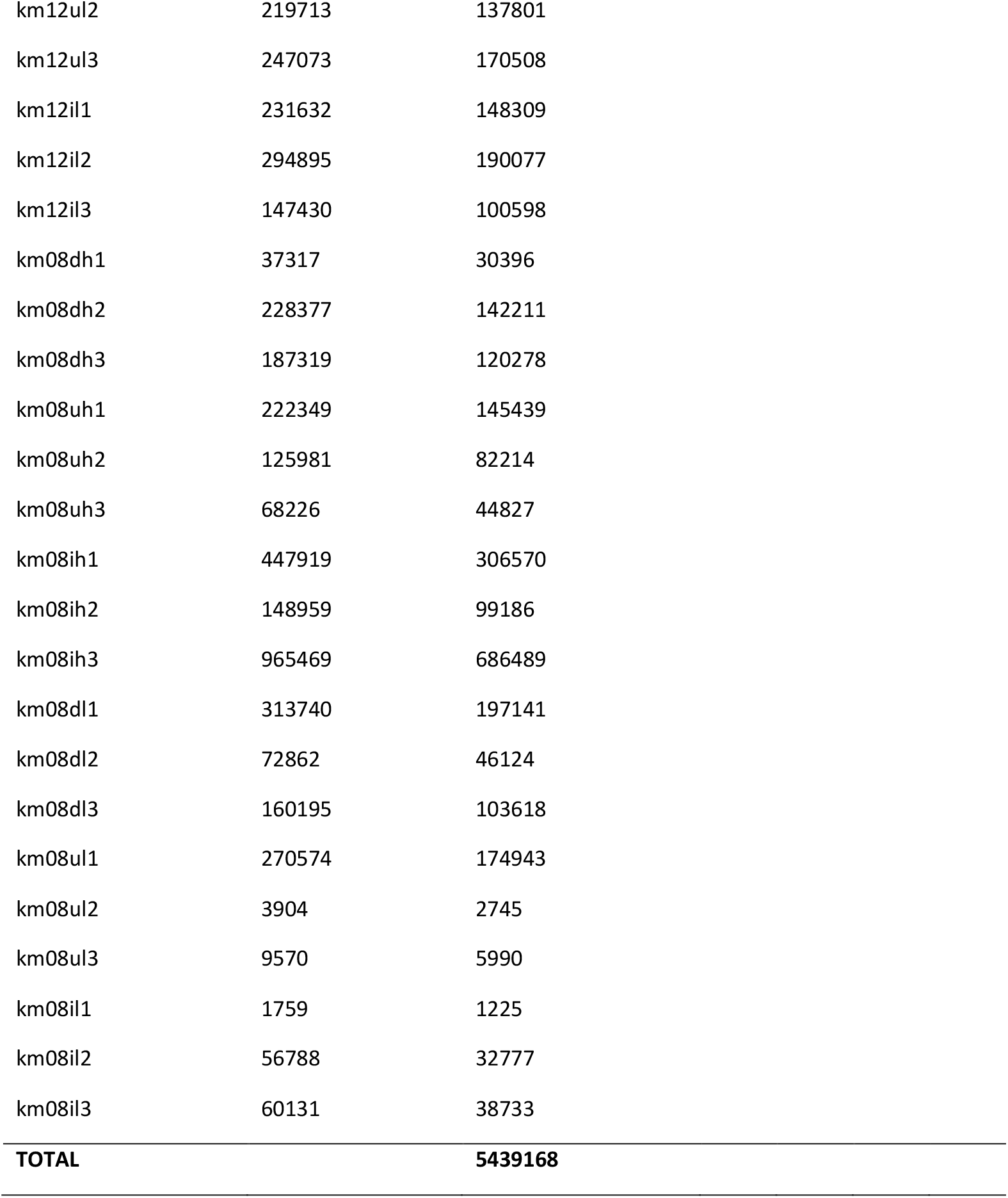
Number of reads filtered DADA2.

**Supplementary Table 2.** Environmental factors (physical) used in the study.

| Sample | Gravel site | point | pH | Temperature | Turbidity | Discharge | EC |
| --- | --- | --- | --- | --- | --- | --- | --- |
| km14dh1 | KM14 | down | 6.5 | 18.4 | 1.8 | 250 | 13.1 |
| km14dh2 | KM14 | down | 6.5 | 18.4 | 1.8 | 250 | 13.1 |
| km14dh3 | KM14 | down | 6.5 | 18.4 | 1.8 | 250 | 13.1 |
| km14ul1 | KM14 | up | 6.5 | 17.8 | 1.2 | 250 | 12.4 |
| km14ul2 | KM14 | up | 7.1 | 17.8 | 1.2 | 250 | 12.4 |
| km14ul3 | KM14 | up | 7.1 | 17.8 | 1.2 | 250 | 12.4 |
| km14dl1 | KM14 | down | 7.1 | 15.7 | 2.2 | 150 | 13.1 |
| km14dl2 | KM14 | down | 7.1 | 15.7 | 2.2 | 150 | 13.1 |
| km14dl3 | KM14 | down | 7.1 | 15.7 | 2.2 | 150 | 13.1 |
| km14uh1 | KM14 | up | 7.1 | 19.1 | 2.2 | 150 | 12.4 |
| km14uh2 | KM14 | up | 7.1 | 19.1 | 2.2 | 150 | 12.4 |
| km14uh3 | KM14 | up | 7.1 | 19.1 | 2.2 | 150 | 12.4 |
| km12dh1 | KM12 | down | 7.5 | 17 | 6.3 | 250 | 13 |
| km12dh2 | KM12 | down | 7.5 | 17 | 6.3 | 250 | 13 |
| km12dh3 | KM12 | down | 7.5 | 17 | 6.3 | 250 | 13 |
| km12uh1 | KM12 | up | 7.5 | 15.6 | 2.8 | 250 | 13.7 |
| km12uh2 | KM12 | up | 7.5 | 15.6 | 2.8 | 250 | 13.7 |
| km12uh3 | KM12 | up | 7.5 | 15.6 | 2.8 | 250 | 13.7 |
| km12ih1 | KM12 | intermediate | 7 | 17 | 2.8 | 250 | 13.7 |
| km12ih2 | KM12 | intermediate | 7 | 17 | 2.8 | 250 | 13.7 |
| km12ih3 | KM12 | intermediate | 7 | 17 | 2.8 | 250 | 13.7 |
| km12dl1 | KM12 | down | 7 | 15.6 | 2.8 | 150 | 13 |
| km12dl2 | KM12 | down | 7 | 15.6 | 2.8 | 150 | 13 |
| km12dl3 | KM12 | down | 7 | 15.6 | 2.8 | 150 | 13 |
| km12ul1 | KM12 | up | 6.9 | 15.6 | 2.8 | 150 | 13.7 |
| km12ul2 | KM12 | up | 6.9 | 15.6 | 2.8 | 150 | 13.7 |
| km12ul3 | KM12 | up | 6.9 | 15.6 | 2.8 | 150 | 13.7 |
| km12il1 | KM12 | intermediate | 7.1 | 15.6 | 2.8 | 150 | 13.7 |
| km12il2 | KM12 | intermediate | 7.1 | 15.6 | 2.8 | 150 | 13.7 |
| km12il3 | KM12 | intermediate | 7.1 | 15.6 | 2.8 | 150 | 13.7 |
| km08dh1 | KM08 | down | 7 | 15.1 | 16.1 | 250 | 11.9 |
| km08dh2 | KM08 | down | 7 | 15.1 | 16.1 | 250 | 11.9 |
| km08dh3 | KM08 | down | 7 | 15.1 | 16.1 | 250 | 11.9 |
| km08uh1 | KM08 | up | 7.1 | 17.7 | 8.1 | 250 | 12.2 |
| km08uh2 | KM08 | up | 7.1 | 17.7 | 8.1 | 250 | 12.2 |
| km08uh3 | KM08 | up | 7.1 | 17.7 | 8.1 | 250 | 12.2 |
| km08ih1 | KM08 | intermediate | 7.9 | 13.8 | 8.3 | 250 | 15 |
| km08ih2 | KM08 | intermediate | 7.9 | 13.8 | 8.3 | 250 | 15 |
| km08ih3 | KM08 | intermediate | 7.9 | 13.8 | 8.3 | 250 | 15 |
| km08dl1 | KM08 | down | 7.1 | 13.8 | 9.1 | 150 | 11.9 |
| km08dl2 | KM08 | down | 7.1 | 17 | 9.1 | 150 | 11.9 |
| km08dl3 | KM08 | down | 7.1 | 17 | 9.1 | 150 | 11.9 |
| km08ul1 | KM08 | up | 7.3 | 17 | 2.2 | 150 | 12.2 |
| km08ul2 | KM08 | up | 7.3 | 15.1 | 2.2 | 150 | 12.2 |
| km08ul3 | KM08 | up | 7.3 | 15.1 | 2.2 | 150 | 12.2 |
| km08il1 | KM08 | intermediate | 7.3 | 15.1 | 6.3 | 150 | 15 |
| km08il2 | KM08 | intermediate | 7.3 | 15.1 | 6.3 | 150 | 15 |
| km08il3 | KM08 | Intermediate | 7.3 | 15.1 | 6.3 | 150 | 15 |

**Supplementary Table 3.** Environmental factors (chemical) used in the study.

| sample | Na | Mg | Al | Ca | Cr | Mn | Fe | Cu | Zn | Cd | Pb | Si | P | F | Cl | NO<sub>3</sub> | SO<sub>4</sub> |
| --- | --- | --- | --- | --- | --- | --- | --- | --- | --- | --- | --- | --- | --- | --- | --- | --- | --- |
| km14dh1 | 5384.1 | 2028.5 | 15.3 | 16594 | 0.6 | 1.28 | 8.61 | 0.5 | 0.6 | 0 | 0 | 5608.9 | 56 | 0.1 | 5 | 2.85 | 9.76 |
| km14dh2 | 5384.1 | 2028.5 | 15.3 | 16594 | 0.6 | 1.28 | 8.61 | 0.5 | 0.6 | 0 | 0 | 5608.9 | 56 | 0.1 | 5 | 2.85 | 9.76 |
| km14dh3 | 5384.1 | 2028.5 | 15.3 | 16594 | 0.6 | 1.28 | 8.61 | 0.5 | 0.6 | 0 | 0 | 5608.9 | 56 | 0.1 | 5 | 2.85 | 9.76 |
| km14ul1 | 5751.5 | 2210.5 | 55.8 | 17234.8 | 0.4 | 3.06 | 5.5 | 0.7 | 6.3 | 0 | 0.2 | 5439.8 | 51.9 | 0.1 | 5 | 2.85 | 9.76 |
| km14ul2 | 5751.5 | 2210.5 | 55.8 | 17234.8 | 0.4 | 3.06 | 5.5 | 0.7 | 6.3 | 0 | 0.2 | 5439.8 | 51.9 | 0.1 | 5 | 2.85 | 9.76 |
| km14ul3 | 5751.5 | 2210.5 | 55.8 | 17234.8 | 0.4 | 3.06 | 5.5 | 0.7 | 6.3 | 0 | 0.2 | 5439.8 | 51.9 | 0.1 | 5 | 2.85 | 9.76 |
| km14dl1 | 5751.5 | 2210.5 | 55.8 | 17234.8 | 0.4 | 3.06 | 5.5 | 0.7 | 6.3 | 0 | 0.2 | 5439.8 | 51.9 | 0.1 | 5 | 2.85 | 9.76 |
| km14dl2 | 5751.5 | 2210.5 | 55.8 | 17234.8 | 0.4 | 3.06 | 5.5 | 0.7 | 6.3 | 0 | 0.2 | 5439.8 | 51.9 | 0.1 | 5 | 2.85 | 9.76 |
| km14dl3 | 5751.5 | 2210.5 | 55.8 | 17234.8 | 0.4 | 3.06 | 5.5 | 0.7 | 6.3 | 0 | 0.2 | 5439.8 | 51.9 | 0.1 | 5 | 2.85 | 9.76 |
| km14uh1 | 5384.1 | 2028.5 | 15.3 | 16594 | 0.6 | 1.28 | 8.61 | 0.5 | 0.6 | 0 | 0 | 5608.9 | 56 | 0.1 | 2 | 2.59 | 9.59 |
| km14uh2 | 5384.1 | 2028.5 | 15.3 | 16594 | 0.6 | 1.28 | 8.61 | 0.5 | 0.6 | 0 | 0 | 5608.9 | 56 | 0.1 | 2 | 2.59 | 9.59 |
| km14uh3 | 5384.1 | 2028.5 | 15.3 | 16594 | 0.6 | 1.28 | 8.61 | 0.5 | 0.6 | 0 | 0 | 5608.9 | 56 | 0.1 | 2 | 2.59 | 9.59 |
| km12dh1 | 5166.9 | 2348.2 | 61.7 | 17379.4 | 0.5 | 4.47 | 57.5 | 0.7 | 0.6 | 0 | 0.1 | 6020.2 | 27.9 | 0.1 | 2 | 2.11 | 9.62 |
| km12dh2 | 5166.9 | 2348.2 | 61.7 | 17379.4 | 0.5 | 4.47 | 57.5 | 0.7 | 0.6 | 0 | 0.1 | 6020.2 | 27.9 | 0.1 | 2 | 2.11 | 9.62 |
| km12dh3 | 5166.9 | 2348.2 | 61.7 | 17379.4 | 0.5 | 4.47 | 57.5 | 0.7 | 0.6 | 0 | 0.1 | 6020.2 | 27.9 | 0.1 | 2 | 2.11 | 9.62 |
| km12uh1 | 5166.9 | 2348.2 | 61.7 | 17379.4 | 0.5 | 4.47 | 57.5 | 0.7 | 0.6 | 0 | 0.1 | 6020.2 | 27.9 | 0.1 | 2 | 2.11 | 9.62 |
| km12uh2 | 5166.9 | 2348.2 | 61.7 | 17379.4 | 0.5 | 4.47 | 57.5 | 0.7 | 0.6 | 0 | 0.1 | 6020.2 | 27.9 | 0.1 | 2 | 2.11 | 9.62 |
| km12uh3 | 5166.9 | 2348.2 | 61.7 | 17379.4 | 0.5 | 4.47 | 57.5 | 0.7 | 0.6 | 0 | 0.1 | 6020.2 | 27.9 | 0.1 | 2 | 2.11 | 9.62 |
| km12ih1 | 5166.9 | 2348.2 | 61.7 | 17379.4 | 0.5 | 4.47 | 57.5 | 0.7 | 0.6 | 0 | 0.1 | 6020.2 | 27.9 | 0.1 | 2 | 2.11 | 9.62 |
| km12ih2 | 5166.9 | 2348.2 | 61.7 | 17379.4 | 0.5 | 4.47 | 57.5 | 0.7 | 0.6 | 0 | 0.1 | 6020.2 | 27.9 | 0.1 | 2 | 2.11 | 9.62 |
| km12ih3 | 5166.9 | 2348.2 | 61.7 | 17379.4 | 0.5 | 4.47 | 57.5 | 0.7 | 0.6 | 0 | 0.1 | 6020.2 | 27.9 | 0.1 | 2 | 2.11 | 9.62 |
| km12dl1 | 5751.5 | 2210.5 | 55.8 | 17234.8 | 0.4 | 3.06 | 5.5 | 0.7 | 6.3 | 0 | 0.2 | 5439.8 | 51.9 | 0.1 | 5 | 2.85 | 9.76 |
| km12dl2 | 5751.5 | 2210.5 | 55.8 | 17234.8 | 0.4 | 3.06 | 5.5 | 0.7 | 6.3 | 0 | 0.2 | 5439.8 | 51.9 | 0.1 | 5 | 2.85 | 9.76 |
| km12dl3 | 5751.5 | 2210.5 | 55.8 | 17234.8 | 0.4 | 3.06 | 5.5 | 0.7 | 6.3 | 0 | 0.2 | 5439.8 | 51.9 | 0.1 | 5 | 2.85 | 9.76 |
| km12ul1 | 5751.5 | 2210.5 | 55.8 | 17234.8 | 0.4 | 3.06 | 5.5 | 0.7 | 6.3 | 0 | 0.2 | 5439.8 | 51.9 | 0.1 | 5 | 2.85 | 9.76 |
| km12ul2 | 5751.5 | 2210.5 | 55.8 | 17234.8 | 0.4 | 3.06 | 5.5 | 0.7 | 6.3 | 0 | 0.2 | 5439.8 | 51.9 | 0.1 | 5 | 2.85 | 9.76 |
| km12ul3 | 5751.5 | 2210.5 | 55.8 | 17234.8 | 0.4 | 3.06 | 5.5 | 0.7 | 6.3 | 0 | 0.2 | 5439.8 | 51.9 | 0.1 | 5 | 2.85 | 9.76 |
| km12il1 | 5751.5 | 2210.5 | 55.8 | 17234.8 | 0.4 | 3.06 | 5.5 | 0.7 | 6.3 | 0 | 0.2 | 5439.8 | 51.9 | 0.1 | 5 | 2.85 | 9.76 |
| km12il2 | 5751.5 | 2210.5 | 55.8 | 17234.8 | 0.4 | 3.06 | 5.5 | 0.7 | 6.3 | 0 | 0.2 | 5439.8 | 51.9 | 0.1 | 5 | 2.85 | 9.76 |
| km12il3 | 5751.5 | 2210.5 | 55.8 | 17234.8 | 0.4 | 3.06 | 5.5 | 0.7 | 6.3 | 0 | 0.2 | 5439.8 | 51.9 | 0.1 | 5 | 2.85 | 9.76 |
| km08dh1 | 4166.4 | 1872 | 75.1 | 14923.4 | 0.5 | 3.72 | 73.6 | 0.6 | 0 | 0 | 0.1 | 5978.8 | 30.3 | 0.1 | 1 | 2.46 | 8.67 |
| km08dh2 | 4166.4 | 1872 | 75.1 | 14923.4 | 0.5 | 3.72 | 73.6 | 0.6 | 0 | 0 | 0.1 | 5978.8 | 30.3 | 0.1 | 1 | 2.46 | 8.67 |
| km08dh3 | 4166.4 | 1872 | 75.1 | 14923.4 | 0.5 | 3.72 | 73.6 | 0.6 | 0 | 0 | 0.1 | 5978.8 | 30.3 | 0.1 | 1 | 2.46 | 8.67 |
| km08uh1 | 3784.3 | 3603.4 | 35.9 | 21083.9 | 0.4 | 7.11 | 37.4 | 0.5 | 3 | 0 | 0.1 | 7440.1 | 24.1 | 0.1 | 1 | 0.35 | 7.82 |
| km08uh2 | 3784.3 | 3603.4 | 35.9 | 21083.9 | 0.4 | 7.11 | 37.4 | 0.5 | 3 | 0 | 0.1 | 7440.1 | 24.1 | 0.1 | 1 | 0.35 | 7.82 |
| km08uh3 | 3784.3 | 3603.4 | 35.9 | 21083.9 | 0.4 | 7.11 | 37.4 | 0.5 | 3 | 0 | 0.1 | 7440.1 | 24.1 | 0.1 | 1 | 0.35 | 7.82 |
| km08ih1 | 5311.5 | 2677.1 | 9.14 | 18844 | 0.4 | 0.56 | 1.75 | 1.4 | 0 | 0 | 0.2 | 6545.8 | 27.3 | 0.1 | 4 | 0.88 | 7.96 |
| km08ih2 | 5311.5 | 2677.1 | 9.14 | 18844 | 0.4 | 0.56 | 1.75 | 1.4 | 0 | 0 | 0.2 | 6545.8 | 27.3 | 0.1 | 4 | 0.88 | 7.96 |
| km08ih3 | 5311.5 | 2677.1 | 9.14 | 18844 | 0.4 | 0.56 | 1.75 | 1.4 | 0 | 0 | 0.2 | 6545.8 | 27.3 | 0.1 | 4 | 0.88 | 7.96 |
| km08dl1 | 4138.3 | 1650.9 | 29.8 | 12388.1 | 0.6 | 19.2 | 39.1 | 1.1 | 3.5 | 0 | 0.2 | 4708.9 | 27.6 | 0.1 | 4 | 2.55 | 9.08 |
| km08dl2 | 4138.3 | 1650.9 | 29.8 | 12388.1 | 0.6 | 19.2 | 39.1 | 1.1 | 3.5 | 0 | 0.2 | 4708.9 | 27.6 | 0.1 | 4 | 2.55 | 9.08 |
| km08dl3 | 4138.3 | 1650.9 | 29.8 | 12388.1 | 0.6 | 19.2 | 39.1 | 1.1 | 3.5 | 0 | 0.2 | 4708.9 | 27.6 | 0.1 | 4 | 2.55 | 9.08 |
| km08ul1 | 4459.1 | 2158.1 | 14.2 | 15607.9 | 0.3 | 3.54 | 0.79 | 0.5 | 5.4 | 0 | 0.1 | 5749.8 | 43.2 | 0.1 | 4 | 2.76 | 8.82 |
| km08ul2 | 4459.1 | 2158.1 | 14.2 | 15607.9 | 0.3 | 3.54 | 0.79 | 0.5 | 5.4 | 0 | 0.1 | 5749.8 | 43.2 | 0.1 | 4 | 2.76 | 8.82 |
| km08ul3 | 4459.1 | 2158.1 | 14.2 | 15607.9 | 0.3 | 3.54 | 0.79 | 0.5 | 5.4 | 0 | 0.1 | 5749.8 | 43.2 | 0.1 | 4 | 2.76 | 8.82 |
| km08il1 | 5311.5 | 2677.1 | 9.14 | 18844 | 0.4 | 0.56 | 1.75 | 1.4 | 0 | 0 | 0.2 | 6545.8 | 27.3 | 0.1 | 4 | 0.88 | 7.96 |
| km08il2 | 5311.5 | 2677.1 | 9.14 | 18844 | 0.4 | 0.56 | 1.75 | 1.4 | 0 | 0 | 0.2 | 6545.8 | 27.3 | 0.1 | 4 | 0.88 | 7.96 |
| km08il3 | 5311.5 | 2677.1 | 9.14 | 18844 | 0.4 | 0.56 | 1.75 | 1.4 | 0 | 0 | 0.2 | 6545.8 | 27.3 | 0.1 | 4 | 0.88 | 7.96 |

**Supplementary Table 4.** Predicted ecological functions of the microbial communities using FAPTOTAX.

In the supplementary excel file

**Supplementary Figure 1.**
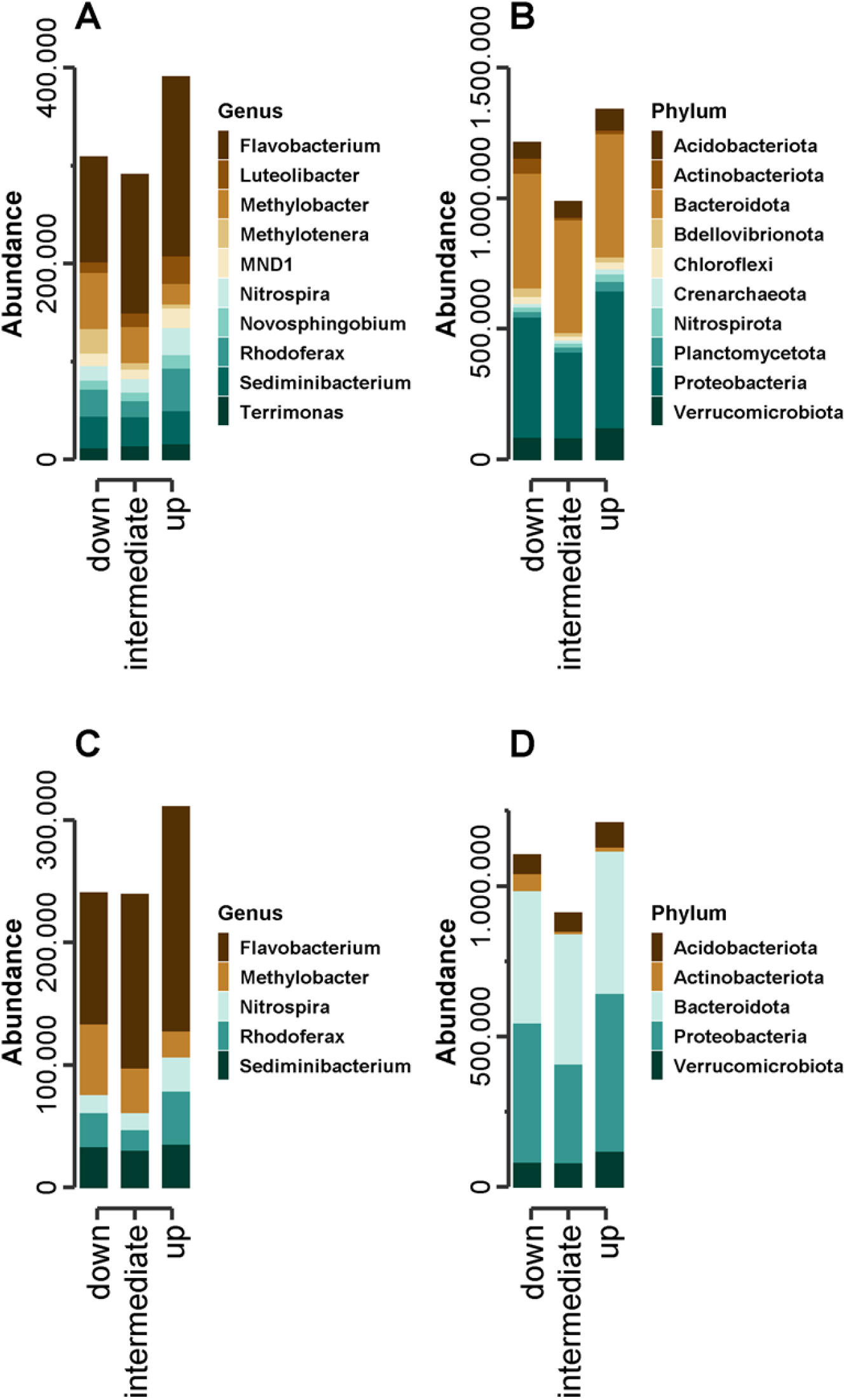
Microbial taxonomy and diversity for each hyporheic point (A) Top 10 most abundant genera. (B) Top 10 most abundant phyla. (C) Top five most abundant genera (D) Top 5 most abundant phyla.

**Supplementary Figure 2.**
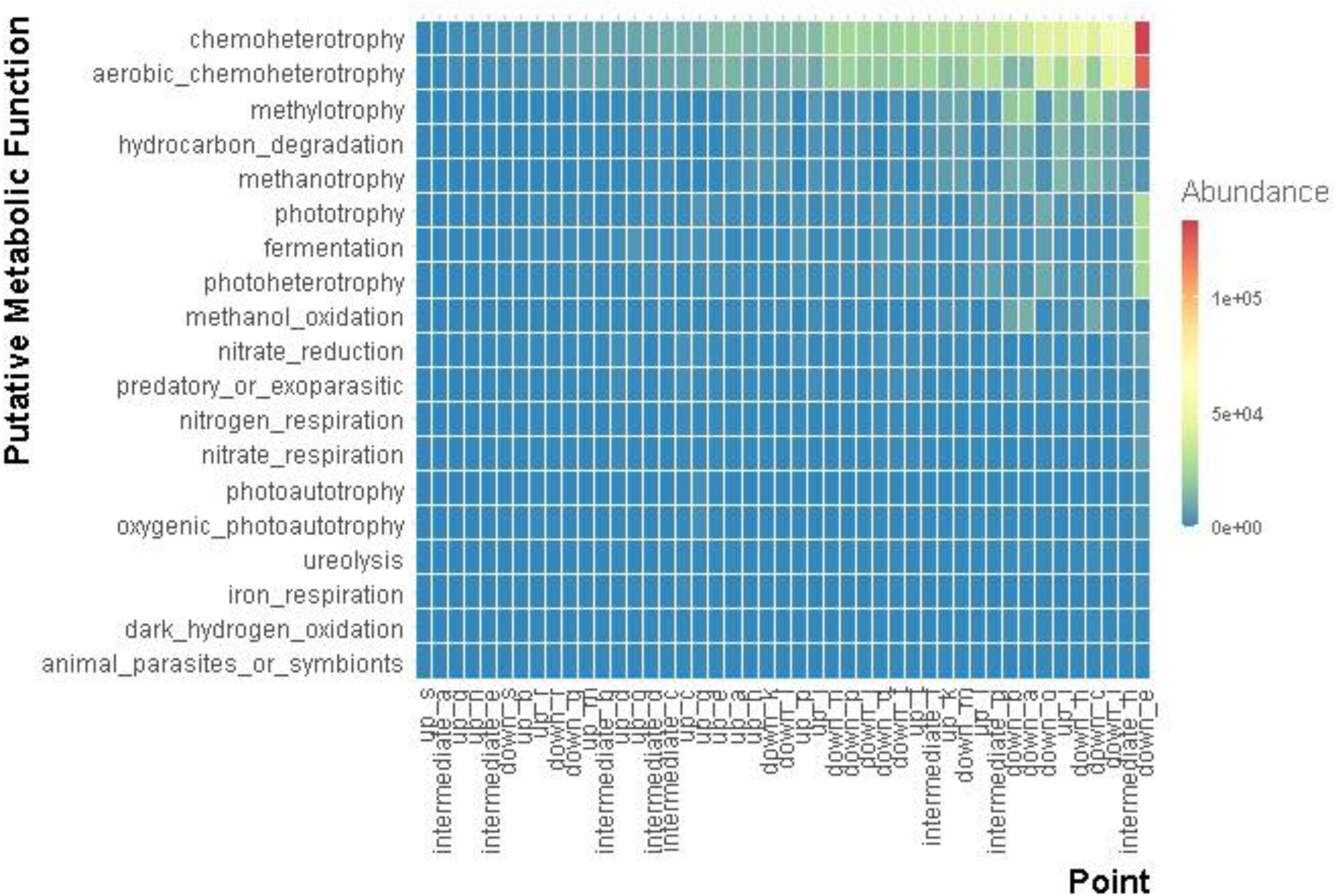
Heatmap showing the abundant putative metabolic function for each hyporheic points i.e., downwelling, upwelling and intermediate.

## Notes

### Competing Interest Statement

The authors have declared no competing interest.

