## Supplementary figures and images for "The influence of river discharge on gravel bar hyporheic microbial community structure and putative metabolic functions"

### supplemental

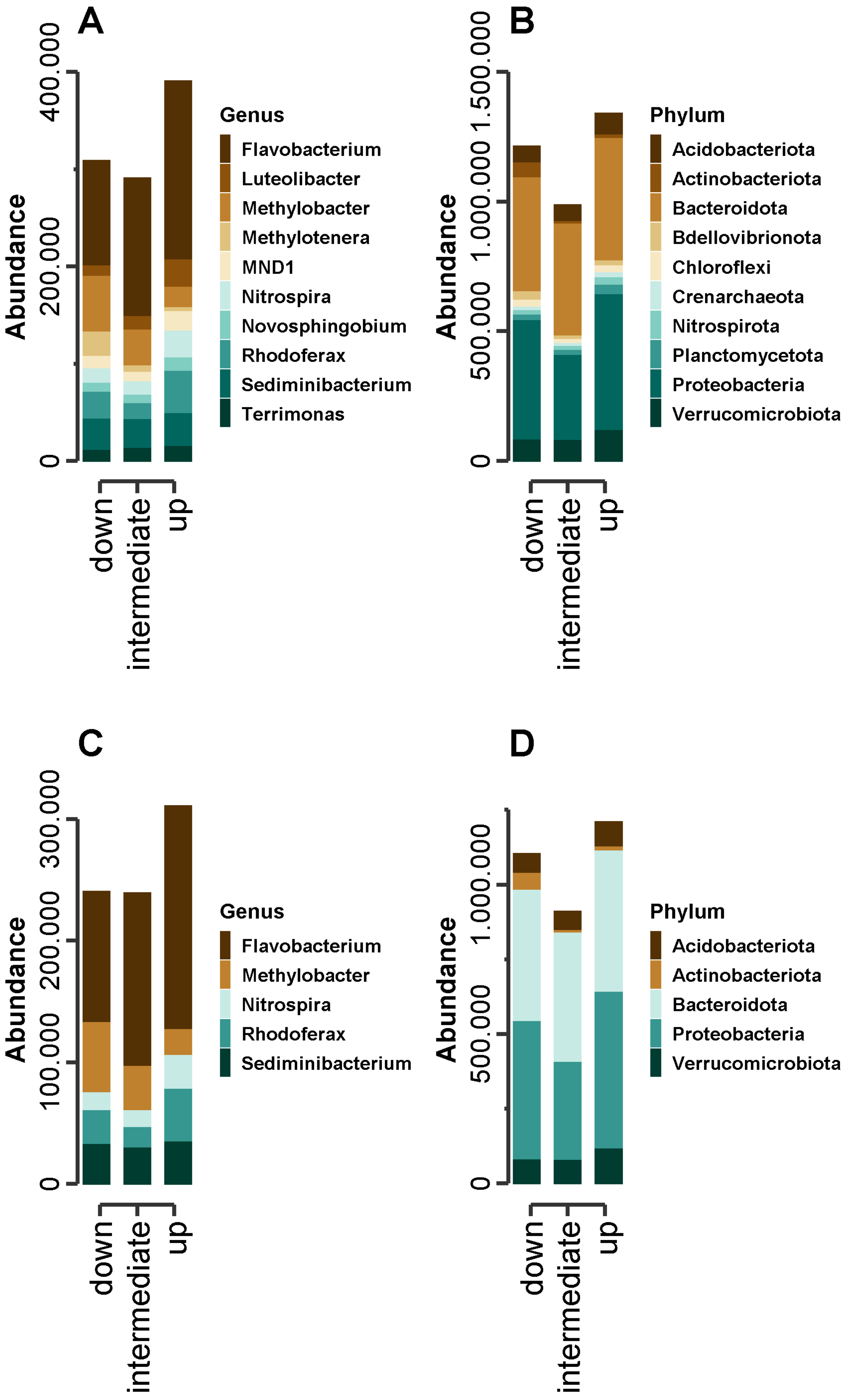

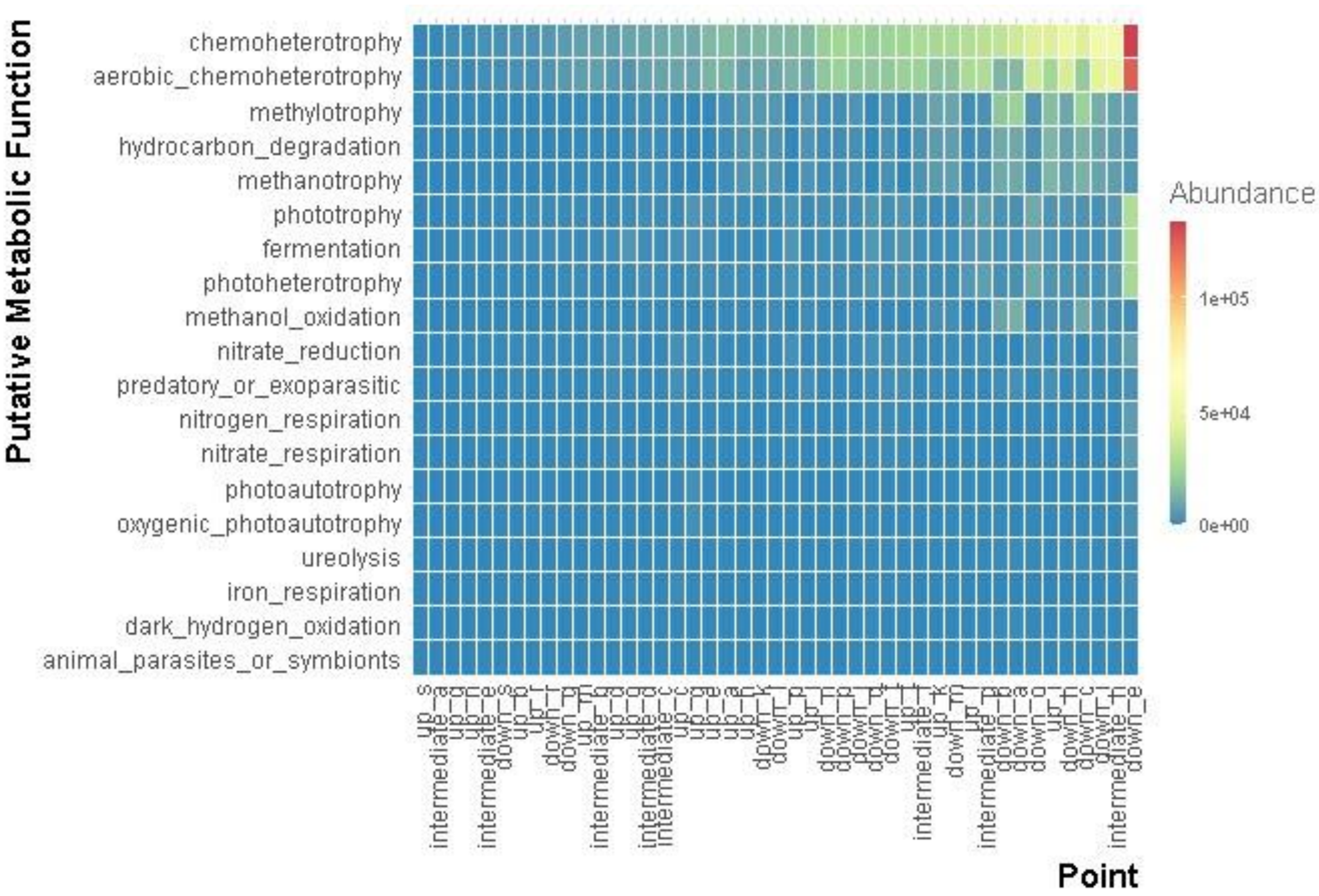
